# The Effects of Different Gait Patterns on Knee Joint Biomechanics and Dynamic Stability during Stair Walking in Healthy Adults

**DOI:** 10.64898/2026.03.19.713073

**Authors:** Gangzheng Yi, Lian Duan, Yuan Gao, Yang Sun, Dapeng Wang

## Abstract

**Objective:** To investigate the effects of different gait patterns on knee joint biomechanics and dynamic stability during stair ascent.

**Methods:** Fourteen healthy males were recruited to ascend stairs using two distinct gait patterns: the “single-step” (leading with the same leg) and “cross-step” (alternating legs) strategies. Kinematic and kinetic data were collected synchronously using a Qualisys infrared motion capture system and a Kistler 3D force plate. Dynamic stability was quantified using the Margin of Stability (MOS), and knee joint biomechanics were evaluated using Patellofemoral Joint Stress (PFJS) and other relevant metrics.

**Results:** Throughout the gait cycle, there was no significant difference in the Medio-Lateral (ML) MOS between the single-step and cross-step patterns (*P*=0.318). However, in the Anterior-Posterior (AP) direction, the MOS for both patterns remained negative and decreased over time, with the cross-step pattern exhibiting significantly lower AP MOS values than the single-step pattern (*P*=0.002). At the moment of left foot-off, significant differences were observed in the right knee joint angle, right knee joint moment, net joint moment, effective quadriceps muscle lever arm, Quadriceps Force (QF), the angle between the quadriceps tendon and patellar ligament, Patellofemoral Joint Force (PFJF), patellofemoral joint stress, and patellofemoral contact area (all *P*<0.001).

**Conclusions:** During stair ascent, the cross-step pattern reduces body stability, thereby increasing the risk of backward falls. Furthermore, this pattern increases patellofemoral joint stress, subjecting the knee to greater loading. Therefore, it is recommended to enhance lower limb muscle strength through targeted training to reduce fall risk. Additionally, adopting a more cautious gait strategy (such as the single-step pattern) can help minimize patellofemoral joint loading and mitigate the risk of patellofemoral pain.

## Introduction

In daily life, stair walking is not only a high-frequency functional activity but also recognized as an efficient form of physical exercise. It has been shown to be more effective than level walking in enhancing lower limb muscle strength, improving cardiorespiratory fitness, controlling body fat, and reducing postprandial hyperglycemia ^[1]^. However, it is also an activity associated with a high risk of falls. Compared to level walking, stair negotiation typically involves greater postural instability, a higher incidence of falls, and distinct lower limb biomechanical characteristics. Consequently, it imposes higher demands on postural control, necessitating greater lower limb joint support moments and knee muscle strength ^[2]^.

Individuals exhibit different gait habits when ascending stairs, primarily categorized as the “single-step” and “cross-step” (or stride) patterns. During single-step ascent, the body maintains center of mass (COM) stability through more frequent lower limb alternations, with the knee and ankle joints performing smaller amplitude flexion-extension and shock absorption movements ^[3]^. In contrast, cross-step ascent requires greater explosive muscle power to propel the body over a larger vertical distance. This results in higher instantaneous loads on the hip and knee joints, accompanied by a significantly increased trunk lean angle and COM displacement range. Different stair ascent patterns directly alter lower limb joint loading, muscle activation states, and balance control strategies, thereby influencing movement efficiency and potential injury risks ^[4]^. Currently, research on stair ascent has largely focused on overall kinematic characteristics or movement optimization in specific populations (such as the elderly ^[5]^or patients with sports injuries ^[6]^), lacking a systematic comparative analysis of the mechanical differences between the fundamental single-step and cross-step gait patterns.

This study aims to compare the differences in dynamic stability and patellofemoral joint mechanics between single-step and cross-step patterns during stair ascent. By investigating how these different gait patterns influence postural control and dynamic stability, this research seeks to enrich the theoretical basis for fall prevention during stair ambulation.

## 1. Participants and Methods

### 1.1 Participants

The sample size was determined using G*Power software. The statistical method was set to an independent samples t-test, with the following parameters: effect size (f) = 0.4, alpha error probability (α) = 0.05, power (1-β) = 0.8, number of groups = 1, number of measurements = 2, and correlation among repeated measures = 0.5. The calculation indicated a minimum required sample size of 11 participants. To account for potential data loss due to motion capture failure, marker displacement, or invalid data, an additional 30% redundancy was added to the sample. Ultimately, a total of 14 healthy males participated in this study. Their basic demographic information is presented below:

**Table 1.** Basic Information of Participants.

| Table 1. Basic Information of Participants |  |  |  |
| --- | --- | --- | --- |
| Age (years) | Height (cm) | Body Mass (kg) | Body Mass Index (kg/m<sup>2</sup>) |
| 19.75±0.86 | 181.24±4.62 | 72.21±7.66 | 22.06±2.94 |

**Inclusion Criteria:** ①No history of neuromusculoskeletal diseases or cognitive impairment, with corrected binocular visual acuity of 1.0 or higher. ②No significant lower limb joint injuries within the past six months. ③No vigorous exercise performed within 24 hours prior to the experiment. ④The right leg was the dominant leg. All subjects were able to understand the experimental procedures and signed an informed consent form.

**Exclusion Criteria:** ①Any gait abnormalities or balance disorders caused by neurological or musculoskeletal injuries. ②A history of lower limb fractures. All subjects participated voluntarily and signed an informed consent form prior to the experiment.

### 1.2 Methods

#### 1.2.1 Instrumentation

A Qualisys three-dimensional (3D) motion capture system (Qualisys, Sweden) consisting of 8 high-speed cameras (Arqus series) was used for kinematic data acquisition and preprocessing at a sampling frequency of 100 Hz. Four Kistler 3D force plates (Model 9260AA6, Kistler Instruments, Switzerland) were embedded into the first, second, third, and fourth steps of a simulated staircase and sampled at 2000 Hz. The force plates were synchronized with the motion capture system via an analog-to-digital converter. Additional equipment included retro-reflective markers, adhesive tape, and kinesiology tape.

#### 1.2.2 Testing Protocol

Prior to the formal experiment, participants were required to change into laboratory-provided tight-fitting clothing and footwear. Subsequently, morphological data were measured, and the dominant leg was determined using a ball-kicking test ^[7]^; all participants were confirmed to have a right dominant leg. Following this, 20 retro-reflective markers were placed on anatomical landmarks of the trunk, pelvis, thighs, shanks, and feet according to the Visual3D model. Before testing, participants performed a 5-minute warm-up and were given sufficient time to familiarize themselves with the experimental environment.

The testing protocol began with a practice trial. Participants stood 15 cm away from the edge of the first step. Upon hearing the “start” command, they initiated movement with the right leg and ascended the stairs at a self-selected comfortable pace until reaching a left-leg single-leg support position to complete the first trial. Participants then returned to the starting position. Three valid trials were recorded for analysis. The experiment included two tasks: ①Single-step pattern: Participants ascended one step at a time. ②Cross-step pattern: Participants ascended two steps at a time.

### 1.3 Data Processing

The stance phase of the dominant leg was extracted for dynamic stability analysis. Data were normalized to 100% of the gait cycle using OriginPro 2024 software. For each participant, data from the three valid trials were averaged for comparative analysis. Based on previous research by Bossé et al. ^[12]^, which identified transitions between different support phases as the most unstable moments during stair negotiation, dynamic stability was analyzed at four critical events:

Right Foot Strike (RFS):Transition from left single-leg support to double support. Left Foot Off (LOFF):Transition from double support to right single-leg support. Left Foot Strike (LFS):Transition from right single-leg support to double support. Right Foot Off (ROFF):Transition from double support to left single-leg support. The foot strike event was defined as the moment when the vertical ground reaction force (vGRF) measured by the force plate reached or exceeded 20 N ^[8]^.

**Figure 1.**
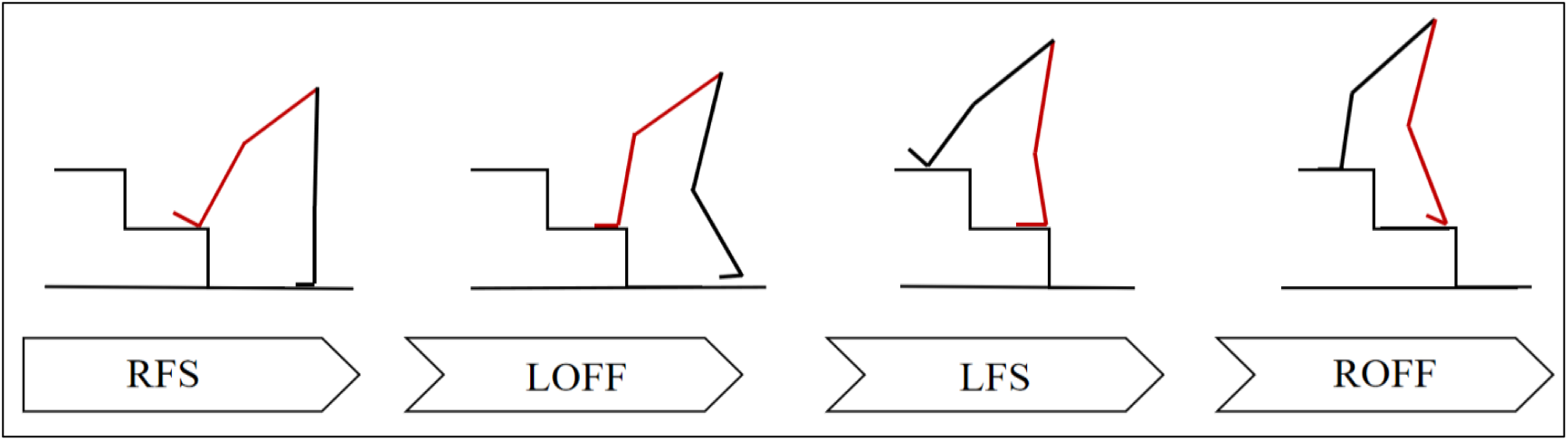
Schematic diagram of the single-step pattern.

**Figure 2.**
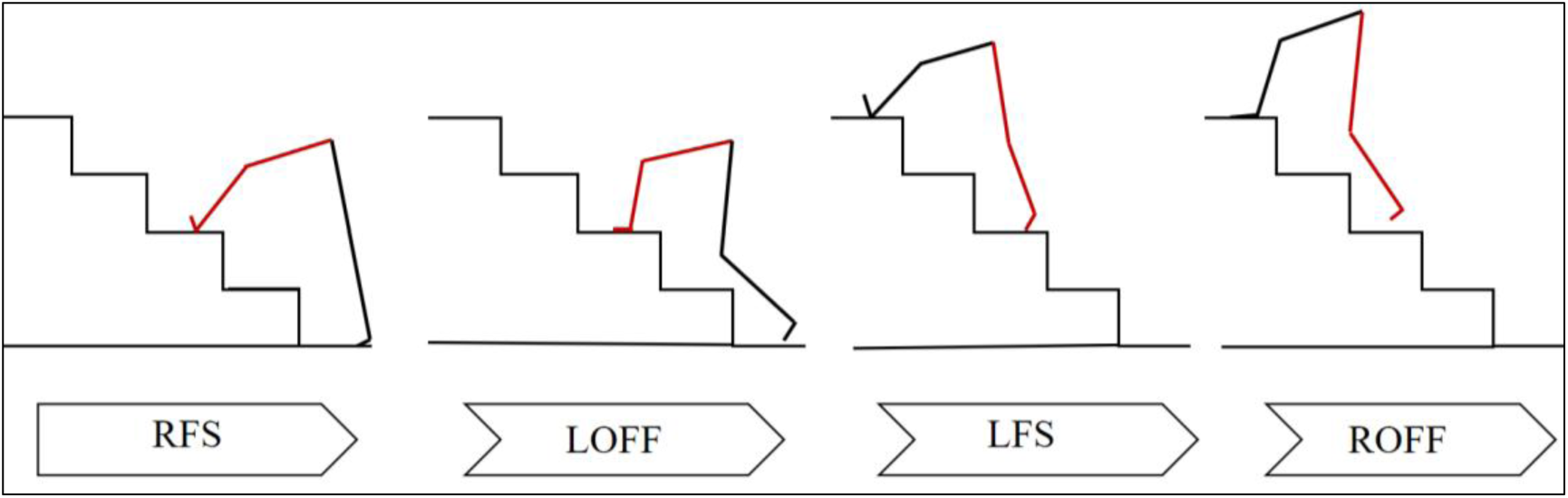
Schematic diagram of the cross-step pattern.

### 1.4 Testing Indicators

#### 1.4.1 Dynamic Stability

Within a complete gait cycle, the Center of Pressure (COP) was defined as the center of the resultant ground reaction force. Step length was defined as the anterior-posterior distance between consecutive heel strikes of the right foot. Step width was defined as the medio-lateral distance between the left and right heels during the double support phase. Step velocity was defined as the time derivative of the displacement of the body’s Center of Mass (COM) in the anterior-posterior direction.

This study analyzed the moments of the dominant leg. The Extrapolated Center of Mass position (XcoM) was defined as the position of the center of mass influenced by velocity. The calculation formulas for the extrapolated center of mass position and dynamic stability are as follows:

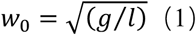

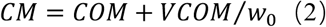

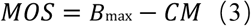

The Margin of Stability (MOS) is based on the inverted pendulum model of the human body. Ground reaction forces act on the anterior-posterior (AP) and medio-lateral (ML) edges of the foot. The body is considered stable when the Extrapolated Center of Mass (XcoM) remains within the boundaries of the Base of Support (BOS). In the equations,ω0represents the natural frequency of the inverted pendulum model, g is the acceleration due to gravity, and l is the length of the inverted pendulum (the vertical distance from the body’s COM to the ground).XCOM(or CM) represents the extrapolated center of mass position, while COMandCOṀ(orVCOM) denote the displacement and velocity of the center of mass at a given moment, respectively. MOSrepresents the dynamic stability at that moment. Bmaxis the maximum boundary of the BOS in a specific direction, which was represented by the position of the Center of Pressure (COP) in this study.

For a given BOS boundary, a positive MOS value indicates that the XcoM lies within the support boundary, signifying body stability. Conversely, a negative MOS value indicates that the XcoM has moved beyond the support boundary, signifying body instability ^[9]^.

In the sagittal plane, a positive value indicates that the COM velocity is directed backward, and the XcoM is positioned anterior to the COP. In the coronal plane, a positive value indicates outward velocity, with the XcoM positioned medial to the COP. Negative values indicate the opposite conditions. A larger absolute MOS value indicates a greater distance between the XcoM and the COP, representing better stability.

**Figure 3.**
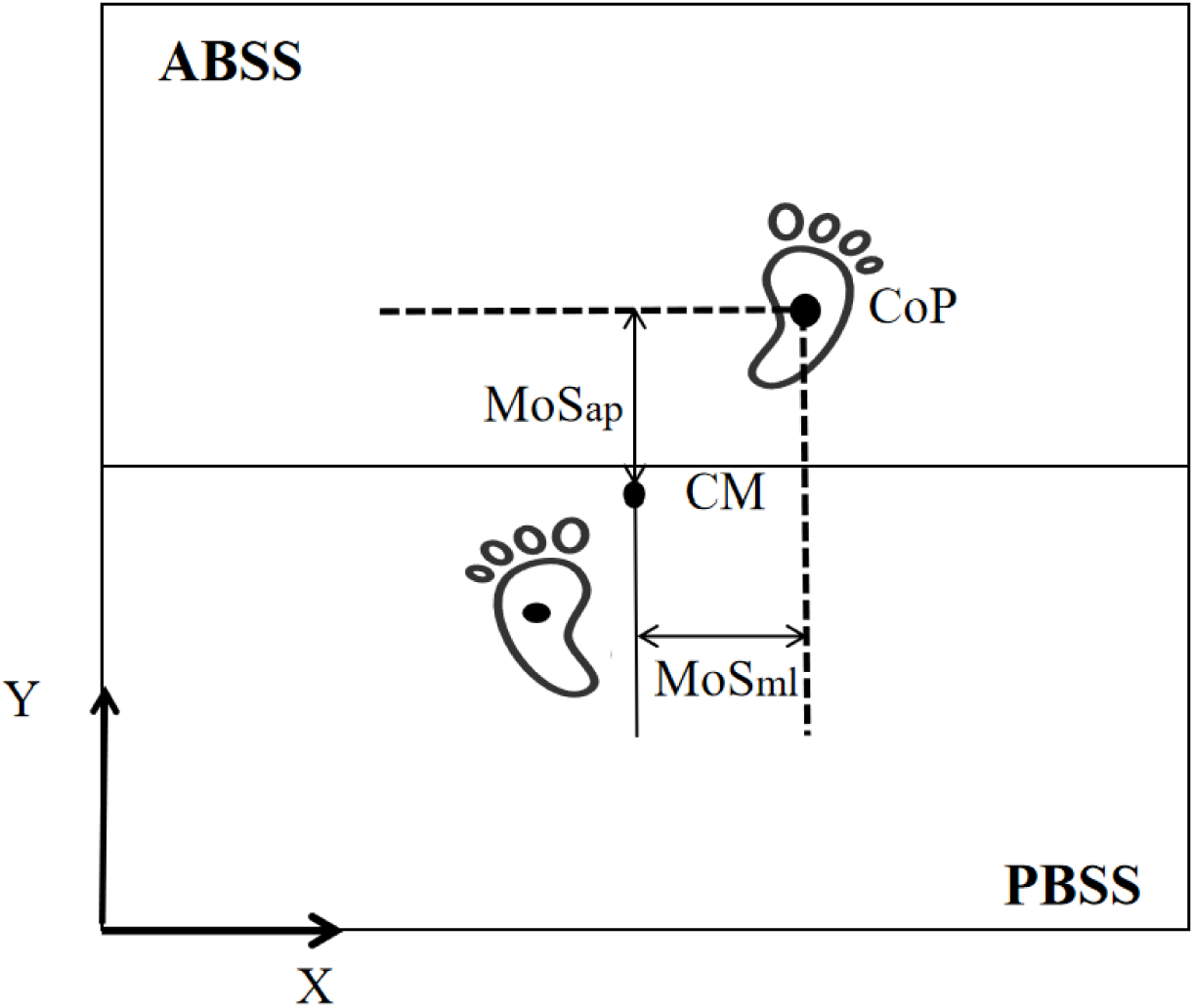
Schematic diagram of dynamic stability. Note: ABSS: Anterior Boundary of the Support Surface.PBSS: Posterior Boundary of the Support Surface

#### 1.4.2 Patellofemoral Joint Stress

Calculation of Patellofemoral Joint Mechanics:

The biomechanical indices of the patellofemoral joint in this study were calculated based on the models proposed by Bresnahan ^[10]^ and Vannatta ^[11]^. The specific calculations are as follows:

Calculation of Quadriceps Force (QF):

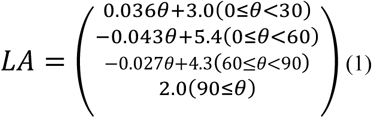

QF ^[10]^ was calculated as the ratio of the knee extension moment to the effective quadriceps moment arm.In Equation (1), the Quadriceps Effective Moment Arm (LA) is defined as a piecewise function of the sagittal plane knee angle (θ).

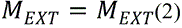

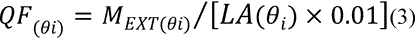

In Equation (2), ***M_EXT_*(*Nm)***(Nm) represents the sagittal plane knee extension moment, and ***M_NET_*(*Nm)***(Nm) represents the sagittal plane net knee joint moment. In Equation (3), θ_*i*_ represents the knee flexion-extension angle at the i-th frame.

Calculation of Patellofemoral Joint Force (PFJF):

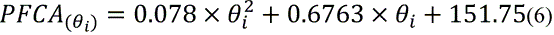

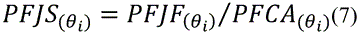

The Patellofemoral Contact Area (PFCA) is a function of the sagittal plane knee angle (θ) and is calculated using Equations (6) and (7). Patellofemoral Joint Stress (PFJS) is defined as the ratio of PFJF to PFCA.

### 1.5 Statistical Analysis

SPSS version 27.0 was used to perform the Shapiro–Wilk test for normality and tests for homogeneity of variance on the dependent variables. Independent samples t-tests were then applied to statistically analyze indicators such as patellofemoral joint stress and dynamic stability. Data are presented as mean ± standard deviation (x±s), and a value of *P*<0.05 was considered statistically significant.

## 2 Results

### 2.1 Differences in Patellofemoral Joint Mechanics Between Different Gait Patterns During Stair Ascent

As illustrated in Figure 4, during stair ascent:

**Figure 4.**
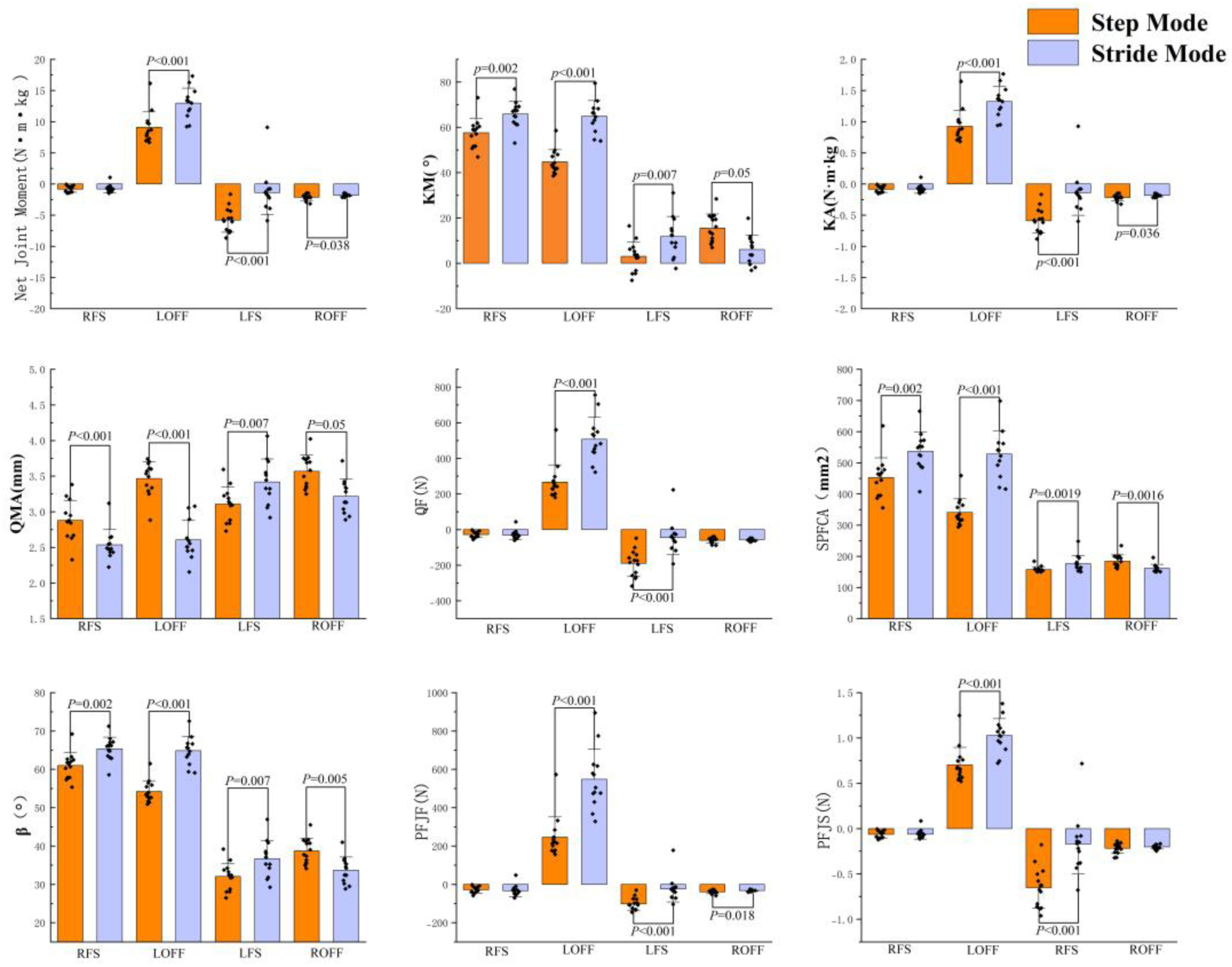
Comparison of biomechanical metrics between single-step and cross-step strategies.

At the Right Foot Strike instant,the stride-over (striding) pattern exhibited significantly greater values for the right knee joint angle (*P*=0.002),the angle between the quadriceps force line and the patellar ligament tension (*P*=0.002),and the patellofemoral contact area (*P*=0.002); conversely,the effective quadriceps moment arm was significantly smaller (*P*<0.001).

At the Left Foot Off instant,the stride-over pattern demonstrated significantly greater values for the right knee joint angle,right knee joint moment,net joint moment,quadriceps muscle force,the angle between the quadriceps force line and the patellar ligament tension,patellofemoral joint force,patellofemoral contact area, and patellofemoral joint stress (all *P*<0.001);the effective quadriceps moment arm was significantly smaller (*P*<0.001).

At the Left Foot Strike instant,the stride-over pattern showed significantly greater values for the right knee joint angle (*P*=0.007),right knee joint moment (*P*<0.001),net joint moment (*P*<0.001),quadriceps muscle force (*P*<0.001),he angle between the quadriceps force line and the patellar ligament tension (*P*=0.007),patellofemoral joint force (*P*<0.001),effective quadriceps moment arm (*P*=0.007),patellofemoral contact area (*P*=0.0019),and patellofemoral joint stress (*P*<0.001).

At the Right Foot Off instant,the stride-over pattern exhibited significantly smaller values for the right knee joint angle (*P*=0.05),effective quadriceps moment arm (*P*=0.05),the angle between the quadriceps force line and the patellar ligament tension (*P*=0.005),and patellofemoral contact area (*P*=0.0016);whereas the right knee joint moment (*P*=0.036),net joint moment (*P*=0.038),and patellofemoral joint force (*P*=0.018) were significantly greater.

### 2.2 Differences in Dynamic Stability Between Gait Patterns During Stair Ascent

As illustrated in Figure 5, at the Right Foot Strike instant, the cross-step strategy exhibited a significantly smaller CMap(*P*=0.002) but a significantly larger VCOMap (*P*<0.001).

**Figure 5.**
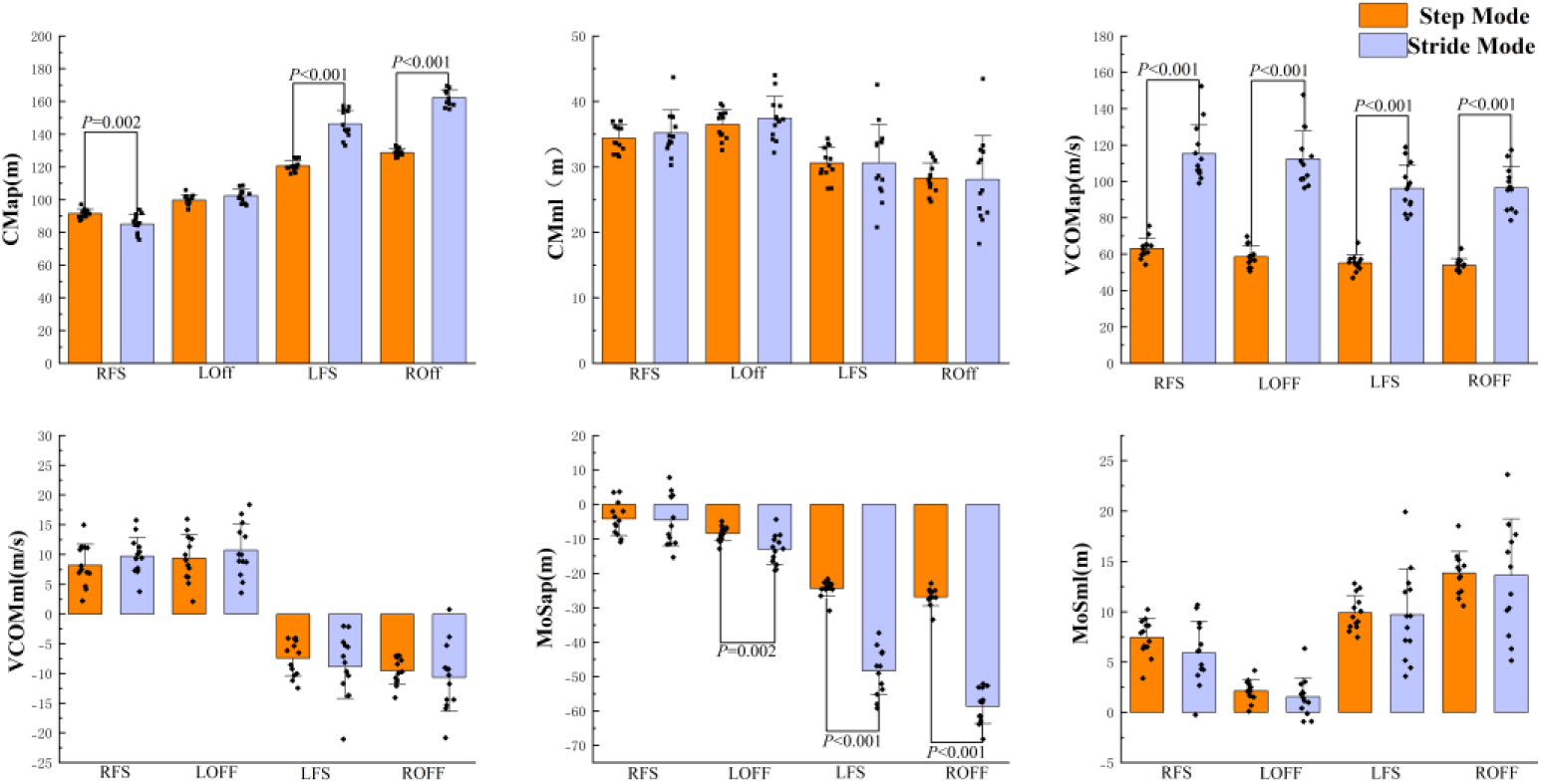
Comparison of dynamic stability between single-step and cross-step strategies. Note:MOS aprepresents the dynamic stability in the anterior-posterior direction,while MOSml represents that in the medial-lateral direction.CMap represents the position of the extrapolated center of mass (XCoM) in the sagittal plane,and CMml represents the position of the extrapolated center of mass in the coronal plane.VCOMaprepresents the component of the center of mass velocity in the sagittal plane, i.e.,the anterior-posterior center of mass velocity;VCOMml represents the component of the center of mass velocity in the coronal plane, i.e., the medial-lateral center of mass velocity.

At the Left Foot Off instant, the cross-step strategy demonstrated a significantly larger VCOMap (*P*<0.001) but a significantly smaller MOSap (*P*=0.002).

At the Left Foot Strike instant, the cross-step strategy showed significantly larger values for both CMap (*P*<0.001) and VCOMap (*P*<0.001),but a significantly smaller MOSap(*P*<0.001).

At the Right Foot Off instant, the cross-step strategy displayed significantly larger values for both CMap (*P*<0.001) and VCOMap (*P*<0.001), but a significantly smaller、MOSap (*P*<0.001).

## 3. Discussion

### 3.1 Differences in Dynamic Stability Between Single-Step and Cross-Step Strategies

The results of this study indicate that, compared with the single-step strategy, the cross-step strategy during stair ascent exhibits a smaller MOSap (Margin of Stability in the anterior-posterior direction), which is negative, suggesting an increased risk of backward falls. However, no significant differences were observed in MOSml (Margin of Stability in the medial-lateral direction) or the factors influencing it. This implies that the cross-step strategy does not demonstrate an elevated risk of falling in the lateral (side-to-side) direction.

Dynamic stability theory is widely regarded as a reliable method for evaluating the human body’s ability to resist perturbations and maintain an upright posture during locomotion.When MOS > 0: The projection of the center of mass (CoM) lies within the boundaries of the Base of Support (BOS). This indicates that the body can effectively adjust its CoM, possesses a stronger ability to maintain an upright posture, and exhibits superior dynamic stability.When MOS < 0: The projection of the CoM falls outside the BOS. This signifies insufficient control over the CoM, making the individual more prone to instability or toppling, thus indicating poor dynamic stability ^[12]^.

The findings of this study reveal that during stair ascent, MOSap undergoes a continuous decrease from negative values. This suggests that MOSap is in an unstable state at the Right Foot Strike instant and gradually transitions into a critically unstable state. Furthermore, the significantly smaller MOSap observed in the cross-step strategy may be attributed to the increased difficulty in controlling the velocity and position of the CoM, leading to poorer dynamic stability in the anterior-posterior direction.

In gait research,dynamic stability refers to the ability to maintain an upright posture and remain stable without falling during walking ^[13][14]^. MOS is a scientific metric that quantifies the real-time relationship among the BOS,CoM position, and velocity to evaluate human dynamic stability^[15][16]^.According to the inverted pendulum model theory, MOS is defined as the difference between Bmax (the boundary of the BOS) and XCoM (Extrapolated Center of Mass). The position and velocity of the body’s CoM play a decisive role in determining the XCoM ^[14]^. For a given direction of support, if the XCoM lies within the BOS, MOS is positive, indicating a stable state; conversely, if the XCoM exceeds Bmax, MOS becomes negative, indicating an unstable state.

Research by Hamel et al.^[17]^ suggests that an increase inVCOM(Velocity of the Center of Mass) is detrimental to controlling the forward displacement of the XCoM,making it difficult to keep the XCoM within the BOS and thereby increasing the risk of backward falls. The present study found that the cross-step strategy during stair ascent is characterized by a larger VCOMap. Based on the calculation method employed in this study,XCoM is a critical factor influencing MOS during stair negotiation. Reducing the XCoM is conducive to improving MOS. However, a larger VCOMap causes the XCoM to increase, thereby decreasing the MOS in the anterior-posterior direction. This finding is consistent with previous research results.

### 3.2 Effects of Single-Step and Cross-Step Strategies on Patellofemoral Joint Biomechanics

During stair ascent, the instant of left foot off serves as a braking phase to resist the downward fall of the center of mass (COM). This phase generates substantial ground reaction forces, braking impulses, and knee extensor moments (KM) ^[19]^. Based on the calculation method of PFJS (Patellofemoral Joint Stress) used in this study, this consequently results in a higher PFJS ^[10]^. At the LOFF instant, body support is primarily provided by the right leg. Furthermore, the right knee exhibits a large flexion angle in the sagittal plane at this point, necessitating greater knee flexion and Quadriceps Femoris (QF) activation to maintain body stability. Therefore, a larger PFJS is also generated at this instant ^[18]^.

Lower limb joint moments serve not only as biomechanical indicators but also as neural control signals, ultimately reflecting the desired control of the central nervous system (CNS). They represent characteristics of neuromuscular control and are considered one of the primary internal factors influencing postural control ability ^[20]^. In the context of stair negotiation, the extension moments of the hip and knee joints, along with the plantarflexion moment of the ankle joint, constitute the core components of the Lower Limb Support Moment. Particularly during stair ascent, the large support moment generated by the lower limbs plays a dual critical role ^[21]^: on one hand, it precisely regulates body posture and ensures dynamic stability; on the other hand, it effectively counteracts the adduction moments induced by trunk and swing leg oscillations. Therefore, the lower limb support moment is a crucial biomechanical factor related to fall risk during stair descent.(Note: The original text mentions “stair descent” in the last sentence of this paragraph; if this is a typo and should refer to “ascent” based on context, please adjust to “ascent”.)

Studies have found that during stair ascent, as the body accelerates forward and upward, the lower limbs require robust support, with knee extension and ankle plantarflexion moments being the primary contributors to the lower limb support moment ^[21]^. The results of this study show that the right knee joint moment peaks at the LOFF instant. At this moment, the right knee joint moment in the cross-step strategy is significantly increased compared to the single-step strategy, ensuring sufficient support for the body during stair ascent and thereby reducing fall risk.

Research by Duncanetal.^[22]^ indicates that stair negotiation leads to high peak patellofemoral joint forces, which is a major cause of patellofemoral pain (PFP). The results of this study demonstrate that patellofemoral joint force increases significantly at the LOFF instant. Furthermore, the cross-step strategy subjects the patellofemoral joint to greater forces and increased loading compared to the single-step strategy. This elevated loading increases the risk of patellofemoral pain. Therefore, it is advisable to adopt a more cautious gait strategy during stair ascent, actively reducing patellofemoral joint stress and loading to mitigate the risk of patellofemoral pain.

## 4 Conclusions

Compared with the single-step strategy, the cross-step strategy during stair ascent results in a significantly smaller MOSap (which is negative), indicating a higher risk of backward falls, and is associated with greater patellofemoral joint stress. The significant increase in peak patellofemoral joint stress may contribute to the development of patellofemoral pain. Therefore, it is recommended to actively adopt a more cautious gait strategy during stair ascent. This can effectively alleviate patellofemoral joint loading and reduce the risk of patellofemoral pain. Additionally, it is suggested to strengthen lower limb training to improve muscle strength, coordination, and control ability, which can effectively lower the risk of falls.

## References

[1] Kang S J, Ahn C H. The effects of home-based stair and normal walking exercises on lower extremity functional ability, fall risk factors, and cardiovascular health risk factors in middle-aged older women[J]. Journal of exercise rehabilitation, 2019, 15(4): 584.

[2] Grimmer M, Zeiss J, Weigand F, et al. Joint power, joint work and lower limb muscle activity for transitions between level walking and stair ambulation at three inclinations[J]. Plos one, 2023, 18(11): e0294161.

[3] Buffinton C M, Blaho R K, Bieryla K A. Biomechanics of single stair climb with implications for inverted pendulum modeling[J]. Journal of Biomechanical Engineering, 2021, 143(8): 081007.

[4] Ko C, Kim J, Choi S, et al. A Comparison on Biomechanical Properties between Step-Over-Step and Step-to-Step Stair Climbing in Young Healthy Male Persons[J]. Available at SSRN 4990663.

[5] Maganaris C, Di Giulio I, Jones D A, et al. Biomechanical and sensory constraints of step and stair negotiation in old age[J]. 2018.

[6] Foot clearance strategy for step-over-step stair climbing in transfemoral amputees

[7] Gribble P A, Tucker W S, White P A. Time-of-day influences on static and dynamic postural control[J]. Journal of athletic training, 2007, 42(1): 35.

[8] Novak A C, Brouwer B. Sagittal and frontal lower limb joint moments during stair ascent and descent in young and older adults[J]. Gait & posture, 2011, 33(1): 54–60.

[9] Havens K L, Mukherjee T, Finley J M. Analysis of biases in dynamic margins of stability introduced by the use of simplified center of mass estimates during walking and turning[J]. Gait & posture, 2018, 59: 162–167.

[10] Bressel E. The influence of ergometer pedaling direction on peak patellofemoral joint forces[J]. Clinical biomechanics, 2001, 16(5): 431–437.

[11] Vannatta C N, Kernozek T W. Patellofemoral joint stress during running with alterations in foot strike pattern[J]. Medicine and science in sports and exercise, 2015, 47(5): 1001–1008.

[12] Bosse I, Oberländer K D, Savelberg H H, et al. Dynamic stability control in younger and older adults during stair descent[J]. Human movement science, 2012, 31(6): 1560–1570.

[13] Hof A L, Gazendam M G J, Sinke W E. The condition for dynamic stability[J]. Journal of biomechanics, 2005, 38(1): 1–8.

[14] Young P M M A, Dingwell J B. Voluntary changes in step width and step length during human walking affect dynamic margins of stability[J]. Gait & posture, 2012, 36(2): 219–224.

[15] Kao P C, Dingwell J B, Higginson J S, et al. Dynamic instability during post-stroke hemiparetic walking[J]. Gait & posture, 2014, 40(3): 457–463.

[16] Mersmann F, Bohm S, Bierbaum S, et al. Young and old adults prioritize dynamic stability control following gait perturbations when performing a concurrent cognitive task[J]. Gait & posture, 2013, 37(3): 373–377.

[17] Hamel K A, Cavanagh P R. Stair performance in people aged 75 and older[J]. Journal of the American Geriatrics Society, 2004, 52(4): 563–567.

[18] Novak A C, Brouwer B. Sagittal and frontal lower limb joint moments during stair ascent and descent in young and older adults[J]. Gait & posture, 2011, 33(1): 54–60.

[19] Reeves N D, Spanjaard M, Mohagheghi A A, et al. Older adults employ alternative strategies to operate within their maximum capabilities when ascending stairs[J]. Journal of Electromyography and Kinesiology, 2009, 19(2): e57–e68.

[20] Reeves ND, Spanjaard M, Mohagheghi AA, et al. Older adults employ alternative strategies to operate within their maximum capabilities when ascending stairs. J Electromyogr Kinesiol. 2009;19(2): e57–e68.

[21] Novak AC, Brouwer B. Sagittal and frontal lower limb joint moments during stair ascent and descent in young and older adults. Gait Posture. 2011;33(1): 54–60.

[22] Duncan R, Peat G, Thomas E, et al. Does isolated patellofemoral osteoarthritis matter?[J]. Osteoarthritis and cartilage, 2009, 17(9): 1151–1155.

